# *ASXL1* mutations that cause Bohring Opitz Syndrome (BOS) or acute myeloid leukemia share epigenomic and transcriptomic signatures

**DOI:** 10.1101/2022.12.15.519823

**Authors:** Isabella Lin, Zain Awamleh, Angela Wei, Bianca Russell, Rosanna Weksberg, Valerie A. Arboleda

## Abstract

*De novo*, truncating variants of *ASXL1* cause two distinct disorders: Bohring-Opitz Syndrome (BOS, OMIM #605039) a rare pediatric disorder characterized by multiorgan anomalies that disrupt normal brain, heart, and bone development causing severe intellectual disability or are somatic driver mutations causing acute myeloid leukemia(AML). Despite their distinct clinical presentations, we propose that *ASXL1* mutations drive common epigenetic and transcriptomic dysregulation in BOS and AML. We analyzed DNA methylation (DNAm) and RNA-seq data from BOS patients (n=13) and controls (n=38) and publicly available DNAm of AML cases with (n=3) and without (n=3) *ASXL1* mutations from The Cancer Genome Atlas (TCGA), and RNA-seq data from AML cases (n=27) from the Beat AML cohort. Using a DNA-methylation based episignature that we previously developed for BOS, we clustered AML, BOS and normal controls together. We showed that AML samples with *ASXL1* mutations clustered closest to individuals with BOS, whereas individuals with AML without *ASXL1* mutations clustered separately. We also observe common dysregulation of the transcriptome between BOS and AML with *ASXL1* mutations compared to controls. Our transcriptomic analysis identified 821 significantly differentially expressed genes that were shared between both data sets and 74.9% showed differential expression in the same direction. BOS patients are rare and have some reports of tumors but no clear guidelines on cancer screening protocols. This represents the first direct comparison between distinct diseases that show common epigenetic and transcriptomic effects, and potentially common drug targets for patients harboring *ASXL1* mutations on the epigenome and transcriptome.

**KEY POINTS:**

- Acute myeloid leukemias harboring somatic *ASXL1* driver mutations and Bohring-Opitz syndrome caused by germline *ASXL1* mutations share common epigenomic and transcriptomic dysregulation
- A gene-centric approach can inform molecular mechanisms across distinct disease types and point towards shared targetable pathways.

---

Despite the significant overlap between genes causing syndromic developmental delay and established cancer driver genes^11^, few studies have established the presence of common, molecular impacts of these mutations across disease-types in patient samples. The initial conceptual links between development and cancer were hypothesized 50 years ago when cancer was considered an “error of development”^2,3^ by Dr. Beatrice Mintz; more specifically, that genetic aberrations in stem cells gave rise to cancer by reversion to an undifferentiated state^3,4^. During human development, genes that regulate the epigenome, called epigenes^5^, control cell-specific RNA expression and complex biological signaling. Mutations involving epigenes are primary genetic drivers in cancer and activate aberrant developmental programs that revert the cells to a malignant stem-cell state^6^. The dual essentiality of epigenes in human development and cancer has been observed over multiple genomic studies^7^, however defining how a pathogenic lesion in the same gene drives epigenomic and molecular mechanisms across distinct diseases has not been shown.

Truncating mutations in the epigene *Additional sex combs-like 1* (*ASXL1*), cause a rare pediatric syndrome, Bohring-Opitz syndrome (BOS, MIM#605309), characterized by intrauterine growth restriction, microcephaly, intellectual disability, dysmorphic features and a characteristic bone-dysmorphic posture. Although growth restricted, BOS patients have an increased risk of Wilms tumor, an embryonic kidney tumor that occurs in children^8^. The same pathogenic *ASXL1* mutations in BOS are somatic driver mutations in 30% of acute myeloid leukemias (AML)^9^. However, no BOS patients have been reported to have developed myelodysplastic syndrome or myeloid leukemias, despite the oldest BOS patient being in their third decade of life. The extent to which the same molecular mechanisms that drive developmental anomalies *in utero* and leukemia somatically remain unclear. To explore whether *ASXL1* mutations dysregulate common epigenetic and transcriptomic targets in development and in AML, we compared the transcriptomic and DNA methylation dysregulation in patients with AML harboring *ASXL1* somatic mutations (AML-ASXL1) and in BOS patients (**Figure 1A**). Our results highlight common dysregulated epigenomic and transcriptomic states driven by protein truncating mutations in *ASXL1*.

**Figure 1.**
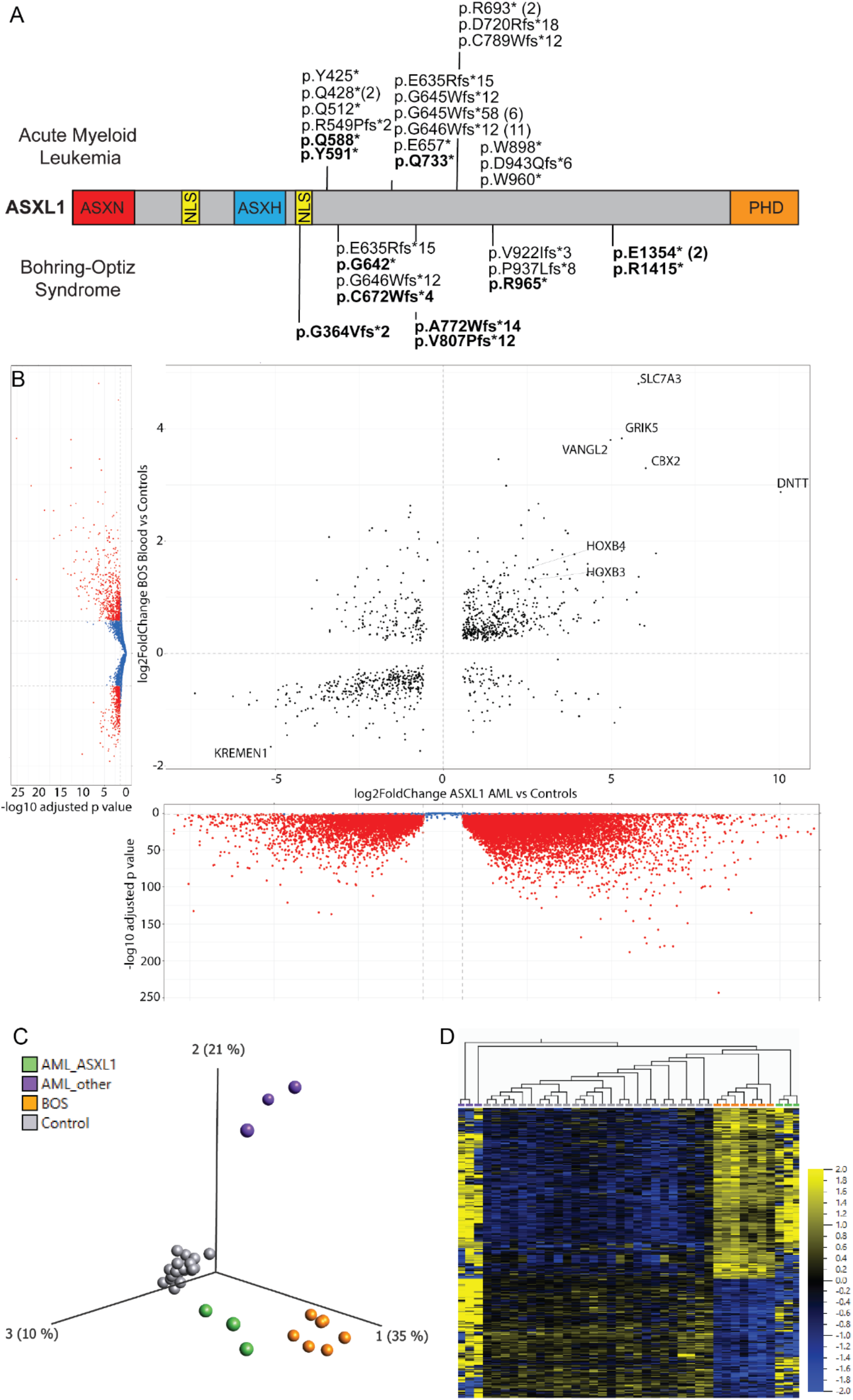
Pathogenic variants in *ASXL1* drive common epigenomic and transcriptional dysregulation in germline disorders and Acute Myeloid Leukemia (AML). **A)** Schematic representation of the ASXL1 protein (GenBank: ASXL1; NM_015338.6; GRCh37), its functional domains, and mutations causing BOS and AML. Red, HB1, ASXL, restriction endonuclease HTH domain (HARE-HTH or ASXN, 11–83); blue, Asx homology domain (ASXH, 236–359); orange, C-terminal plant homeodomain (PHD, 1506–1539). Mutations in bold have both RNA-seq and DNA methylation data. **B)** Global transcriptomic analysis of BOS-patient whole blood compared to controls identified 2118 p-adj significant genes (blue and red) and 1097 differentially expressed genes (DEGs, red) with absolute log fold change greater than 1.5 (vertical axis). Transcriptomic analysis comparing *ASXL1*-mutated AML with samples without *ASXL1* mutations identified 23,943 DEGs (horizontal axis). DEGs are Bonferroni-corrected with p-value, p < 0.05. DEGs with a log2 fold change > 0.58, corresponding to an absolute fold change greater than 1.5 are shown in red). Our study identified 821 genes that were differentially expressed across all ASXL1 mutated samples, regardless of sample or disease types. **C)** principal component analysis (PCA) plot and **D)** heatmap showing clustering of individuals with BOS (n=8; orange), typically developing controls (n=26; grey), individuals with AML caused by a somatic mutation in *ASXL1* (n=3; green), and individuals with AML caused by somatic mutations in other genes (n=3; purple). Clustering is based on DNA methylation values for each individual at 413/763 CpG sites identified in the BOS DNAm signature. The heatmap color gradient indicates the normalized DNAm value ranging from -2.0 (blue) to 2.0 (yellow). Euclidean distance metric is used in the heatmap clustering dendrograms.

Informed consent was obtained from all research participants according to the protocol approved by the Hospital for Sick Children (REB#1000038847) and UCLA (IRB#11-001087). DNA was extracted from blood and processed as previously described^10^. RNA was extracted from PAXgene Blood Tubes (Qiagen) using the MagMAX™ RNA Isolation Kit (Ambion) and libraries were prepared using Truseq Stranded TotalRNA LibraryPrep Gold (Illumina) with QiaSelect rRNA and globin depletion (Qiagen). Libraries were pooled and sequenced to 40 million reads per sample on a NovaSeq6000. We compared transcriptomic data from 27 AML-ASXL1 samples from the Beat AML cohort^11^ with non-AML blood controls. All RNA-seq data was processed through our RNA-seq pipeline where reads are mapped to hg38 using STAR 2.7.0e^12^. Gene counts from raw reads were generated using featureCounts1.6.5 and differential expression was quantified using DESeq2v1.24.0^13^, corrected for batch, sex, age and disease state (Table S1 and S2).

Blood transcriptomic analysis comparing BOS to controls (**Figure 1B**, vertical) identified 2,118 significant differentially expressed genes (DEGs) (p_adj_ < 0.05), with 1097 DEGs with an absolute fold change greater than 1.5. Parallel analysis comparing AML-ASXL1 patient samples compared with control blood (**Figure 1B**, horizontal) identified 23,976 DEGs (11,394 protein coding DEGs), likely due to the significant cell-type heterogeneity in AML specimens. To identify DEGs that are common across different disease-types driven by *ASXL1*-mutations, we plotted fold change for the transcriptomic analyses across their respective axes (**Figure 1B, center**) which highlighted 821 significant genes with an absolute fold change greater than 1.5. Most of the transcriptomic dysregulation occurs in the same direction (615/821, 74.9%) with 53.1% (327/615) of DEGs showing increased gene expression associated with presence of ASXL1 mutations. The intersection of these gene lists was shown to be significant by fisher exact test (p-value= 2.059301e-90). Our data also highlights genes identified in our previous studies comparing transcriptomic dysregulation across different cell types in BOS^14^. These genes include *VANGL2*, a member of the planar cell polarity pathway^15^ and *GRIK5*, a pre- and post-synaptic receptor for glutamate, a crucial excitatory neurotransmitter of the central nervous system^14,16^.

We had previously established a BOS-specific DNA methylation episignature^10^ that used 763 sites to distinguish pathogenic ASXL1 mutations from normotypic matched controls and from patients harboring variants of uncertain significance in *ASXL1*. To investigate whether AML-ASXL1 samples and BOS samples share epigenetic mechanisms, we obtained Illumina 450K DNAm data for six AML samples from The Cancer Genome Atlas (TCGA)^17^ data on the Genomic Data Commons (GDC) repository^18^. Of these, three individuals harbored a somatic mutation in *ASXL1*, while the remaining three harbored somatic mutations in other genes (AML control). We compared DNAm episignature profiles of typically developing controls (grey, n=26), individuals with BOS (orange, n=8), AML-ASXL1 samples (green, n=3) and AML controls (purple, n=3). The principal component analysis (PCA, **Figure 1C**) and hierarchical clustering (dendrogram, **Figure 1D**) using the BOS DNAm episignature sites (413 CpG sites available for comparison)^10^ showed the AML-*ASXL1* individuals (green) clearly distinguished from controls on the first principal component (35%) and from individuals with BOS (orange) on the third principal component (10%) (**Figure 1C**). The three AML controls (purple) are also distinguished from both controls and individuals with BOS on the second principal component (21%). When performing unsupervised clustering of all samples, AML-A*SXL1* samples clustered more closely with individuals with BOS, whereas AML controls clustered separately. This clustering pattern was recapitulated in the dendrogram and heat map (**Figure 1D**), which showed greater overlap in methylation patterns between individuals with BOS (orange) and AML-*ASXL1* (green) compared to AML control (purple). The mean methylation value for the four groups at each of the 413 BOS episignature sites and Cpg site annotations are in Table S3. These findings demonstrate that the DNA methylation signal associated with *ASXL1* mutations supersedes the DNA methylation signals associated with disease status (BOS vs AML), age, and sex.

Our integrative analysis demonstrates that truncating mutations in *ASXL1* drive a common profile of molecular dysregulation, regardless of disease-type or origin (germline versus somatic) of the *ASXL1* mutation. The cell-type and developmental contexts remain critical to determining the complete spectrum of pathological effects of *ASXL1* mutations, as *ASXL1* is known to act as part of protein complexes which require co-expression of other complex factors^19^. We have recently shown that BOS patients’ cells have overactive Wnt signaling^14^ which may drive the increased risk of Wilms tumor^8,20,21^. Interestingly, there are no reported myeloid malignancies in BOS, which may be a product of the *ASXL1* mutation existing in both microenvironment stromal cells and the hematopoietic stem cells, rather than in just the cancer-prone HSC^22^. Comparative studies between germline syndromes and malignancies driven by the same gene can identify novel mechanisms and inform repurposing of existing drugs for orphan rare diseases^23^. These data highlight the potential for emerging epigenetic therapies designed for cancer to be repurposed in rare pediatric syndromes driven by shared mutations and mechanisms^24^.

## Public Data Sets

https://portal.gdc.cancer.gov/projects/TCGA-LAML.

All relevant raw transcriptomic data in the generation of figure one is available through the GEO under accession number (TBD). The DNA methylation datasets generated during the current study are not publicly available due to institutional ethical restrictions but are available from the corresponding author on reasonable request to authors.

## Author Contributions

VA, ZA and IL designed and conceptualized the study and wrote the paper. BR coordinated sample collection for patient samples. IL, SL and AW performed data generation and transcriptomic analysis for patient-derived samples. ZA and RW performed data generation and analysis of DNA methylation data. All authors contributed to the writing of the manuscript.

## Acknowledgements

We would like to thank Seung Hyuk T. Lee who initiated early aspects of this work. We would also like to thank the patients and families who contributed to our ASXL1 biobank which allowed us to ask these questions across many sample types. Finally, we would like to thank the BEAT-AML cohort team for making their data publicly available to answer important questions.

## REFERENCES

1. Qi H, Dong C, Chung WK, Wang K, Shen Y. Deep genetic connection between cancer and developmental disorders. Hum. Mutat. 2016;37(10):1042–1050.

2. Mintz B. Gene expression in neoplasia and differentiation. Harvey Lect. 1978;71:193–246.

3. Bellacosa A. Developmental disease and cancer: biological and clinical overlaps. Am. J. Med. Genet. A. 2013;161A(11):2788–2796.

4. Shipitsin M, Polyak K. The cancer stem cell hypothesis: in search of definitions, markers, and relevance. Lab. Invest. 2008;88(5):459–463.

5. Medvedeva YA, Lennartsson A, Ehsani R, et al. EpiFactors: a comprehensive database of human epigenetic factors and complexes. Database. 2015;2015:bav067.

6. Fahrner JA, Bjornsson HT. Mendelian disorders of the epigenetic machinery: tipping the balance of chromatin states. Annu. Rev. Genomics Hum. Genet. 2014;15:269–293.

7. Nussinov R, Tsai C-J, Jang H. How can same-gene mutations promote both cancer and developmental disorders? Sci Adv. 2022;8(2):eabm2059.

8. Russell B, Johnston JJ, Biesecker LG, et al. Clinical management of patients with ASXL1 mutations and Bohring-Opitz syndrome, emphasizing the need for Wilms tumor surveillance. Am. J. Med. Genet. A. 2015;167A(9):2122–2131.

9. Micol J-B, Abdel-Wahab O. The Role of Additional Sex Combs-Like Proteins in Cancer. Cold Spring Harb. Perspect. Med. 2016;6(10.):

10. Awamleh Z, Chater-Diehl E, Choufani S, et al. DNA methylation signature associated with Bohring-Opitz syndrome: a new tool for functional classification of variants in ASXL genes. Eur. J. Hum. Genet. 2022;

11. Tyner JW, Tognon CE, Bottomly D, et al. Functional genomic landscape of acute myeloid leukaemia. Nature. 2018;562(7728):526–531.

12. Dobin A, Gingeras TR. Optimizing RNA-Seq Mapping with STAR. Methods Mol. Biol. 2016;1415:245–262.

13. Yabumoto M, Kianmahd J, Singh M, et al. Novel variants in KAT6B spectrum of disorders expand our knowledge of clinical manifestations and molecular mechanisms. Mol Genet Genomic Med. 2021;9(10):e1809.

14. Lin I, Wei A, Awamleh Z, et al. Multi-omics on truncating ASXL1 mutations in Bohring Opitz syndrome identify dysregulation of canonical and non-canonical Wnt signaling. bioRxiv. 2022;2022.12.15.520167.

15. Galea GL, Maniou E, Edwards TJ, et al. Cell non-autonomy amplifies disruption of neurulation by mosaic Vangl2 deletion in mice. Nat. Commun. 2021;12(1):1159.

16. Chew LJ, Yuan X, Scherer SE, et al. Characterization of the rat GRIK5 kainate receptor subunit gene promoter and its intragenic regions involved in neural cell specificity. J. Biol. Chem. 2001;276(45):42162–42171.

17. Cancer Genome Atlas Research Network, Weinstein JN, Collisson EA, et al. The Cancer Genome Atlas Pan-Cancer analysis project. Nat. Genet. 2013;45(10):1113–1120.

18. Jensen MA, Ferretti V, Grossman RL, Staudt LM. The NCI Genomic Data Commons as an engine for precision medicine. Blood. 2017;130(4):453–459.

19. Katoh M. Functional and cancer genomics of ASXL family members. Br. J. Cancer. 2013;109(2):299–306.

20. Carraro DM, Ramalho RF, Maschietto M. Gene Expression in Wilms Tumor: Disturbance of the Wnt Signaling Pathway and MicroRNA Biogenesis. Codon Publications; 2016.

21. Corbin M, de Reyniès A, Rickman DS, et al. WNT/beta-catenin pathway activation in Wilms tumors: a unifying mechanism with multiple entries? Genes Chromosomes Cancer. 2009;48(9):816–827.

22. Zhang P, Chen Z, Li R, et al. Loss of ASXL1 in the bone marrow niche dysregulates hematopoietic stem and progenitor cell fates. Cell Discov. 2018;4:4.

23. Roessler HI, Knoers NVAM, van Haelst MM, van Haaften G. Drug Repurposing for Rare Diseases. Trends Pharmacol. Sci. 2021;42(4):255–267.

24. Cheng Y, He C, Wang M, et al. Targeting epigenetic regulators for cancer therapy: mechanisms and advances in clinical trials. Signal Transduction and Targeted Therapy. 2019;4(1):1–39.

